# Population Structure of *Leishmania* using POPSICLE reveals extensive dichotomy in zygosity and discloses the role of sex in diversity of the parasite

**DOI:** 10.1101/420323

**Authors:** Jahangheer S. Shaik

## Abstract

Mosaic aneuploidy prevalent in organisms such as *Leishmania* and *Fungi* and in genomes of some neurological disorders and cancers manifest as non-integer haplotypes due to heterogeneity in somy across a population of cells. Thus, the tools designed for strictly haploid or diploid genomes are insufficient to study them. We addressed this issue by upgrading our population genetics tool POPSICLE for aneuploid genomes and studied the population structure of 50 strains of *Leishmania* to understand genetic diversity and the sexual strategies that predispose to that diversity. *Leishmania* showed enormous diversity but a dichotomic nature of extreme zygosities. To understand this dichotomy, we specifically studied two species, *L. tropica* that contained strains with both hetero and homozygosities and *L. major* that were mainly homozygous. The homozygosity in *L. tropica* was a consequence of extreme inbreeding while heterozygosity was due to recent hybridizations involving two different genotypes. In contrast, *L. major* also contained two different genotypes and products of extreme inbreeding but no recent hybridizations. The heterozygous strains of *L. tropica* that were geographically isolated from the homozygous strains were F1 hybrids that appeared sterile to each other while those in proximity to the homozygous strains were outcrosses involving multiple cycles of hybridization indicating their mating preference with homozygous strains. Development of POPSICLE for aneuploid genomes offers a unique tool for determining the shared ancestries and in reinforcing sex as one of the driving mechanisms for speciation as demonstrated for *Leishmania*. POPSICLE is a Java based utility available for free download at https://popsicle-admixture.sourceforge.io/

## Introduction

Protozoan parasites of the genus *Leishmania* produce a spectrum of diseases such as localized cutaneous lesions, chronic and destructive mucocutaneous infections and lethal visceral infections. Leishmaniasis is endemic in 98 different countries with an estimated 1.5 million cases per year, an incidence of 1 million of new cases per year (WHO. 2018) and 350 million more at the risk (Murray et al. 2005; Alvar et al. 2012). Leishmaniasis is the second major cause of parasite-related death after malaria with approximately 20-30,000 deaths per year (Desjeux 1992; Alvar et al. 2012). There are currently 53 recognized species, of which at least 20, mainly of the subgenra *Leishmania* and *Viannia*, infect humans (Jackson et al. 1986; Kevric et al. 2015). *Leishmania* have very diverse biology, morphology, vector usage, vertebrate reservoirs and differences in disease tropism (Atayde et al. 2014; Brettmann et al. 2016). Despite this observed diversity, at a molecular level their genomes are highly syntenic and contain a large proportion of conserved genes (Peacock et al. 2008). The observed diversity could be dictated by epigenomic control, the presence of a few species-specific genes or by aneuploidy to control the mRNA levels (Beverley 1991; Sterkers et al. 2012; Sterkers et al. 2014), (Dumetz et al. 2017; Inbar et al. 2017). Other factors such as gradual accumulation of mutations during a clonal expansion or through sexual recombination producing admixtures of divergent genomes remain a matter of considerable debate (Bastein et al. 1992; Tibayrenc et al. 1993; Banuls et al. 1997b; Ravel et al. 2006a; Nolder et al. 2007b; Rougeron et al. 2009; Rougeron et al. 2010).

The isolation of *Leishmania* strains that have been characterized as putative hybrids is now well described (Kato et al. 2016), and the panmictic generation of experimental hybrids, where there is no barrier to intra or inter-species mating, has been formally demonstrated in a laboratory setting (Inbar et al. 2013; Romano et al. 2014). Multi locus genotyping using techniques such as multilocus enzyme electrophoresis (MLEE), random amplified polymorphic DNA (RAPD), and multilocus microsatellite typing (MLMT) have suggested hybrids between closely related new world species (Belli et al. 1994; Dujardin et al. 1995; Banuls et al. 1997a; Torrico et al. 1999; Nolder et al. 2007a), between closely related old-world species (Evans et al. 1987; Kelly et al. 1991; Odiwuor et al. 2011), and between two extremely divergent species, *L. infantum* and *L. major* (Ravel et al. 2006b). Using more discriminatory genotyping approaches such as MLMT, and especially whole genome sequencing (WGS), natural hybridization has also been reported at the intraspecies level for *L. infantum, L. donovani*, and *L. tropica* (Chargui et al. 2009; Rogers et al. 2014a; Iantorno et al. 2017). Therefore, the debate regarding *Leishmania* reproductive strategies mainly concerns the mode of genetic exchange and its impact on population structure, not whether it occurs. Although the studies addressing divergence within species have previously been conducted, a comprehensive cross-species analysis using genome-wide markers is long overdue (Peacock et al. 2008; Valdivia et al. 2015). The revealing of shared ancestries argues strongly in favor of sexual recombination and its positive role in speciation of this organism, however the employment of tools designed for strictly haploid and diploid genomes are insufficient for *Leishmania* and may lead to the drawing improper conclusions. Thus, we designed a novel population genetics tool called POPSICLE to gain insight into the sexual strategies in *Leishmania*.

Population genetics is a field that offers strategies to understand the mode of sex in organisms. Some of the strategies include evaluating linkage disequilibrium, heterozygosity because of haplotypes inherited from different genotypes and finding shared ancestries across populations which are otherwise considered different by strain, species, or genus classifications. When used independently, these strategies may sometimes provide ambiguous results. For example, strong linkage disequilibrium has traditionally been linked to clonality. However, extreme inbreeding generates homozygosity that can be easily argued to be clonality (Tibayrenc et al. 1993; Rougeron et al. 2010). Thus, multiple strategies leading to similar conclusions when independently employed or when supplemented with each other, may reduce Type-I and Type-II errors, leading to higher confidence in the conclusions reached. Another factor to consider is that the tools employed should be relevant for the genomes being studied. For example, ancestry determining tools such as structure (Pritchard et al. 2000) and fine-structure (Lawson et al. 2012) can be readily applied for haploid genomes and can be used for diploid genomes post haplotype phasing. They are not relevant for *Leishmania* due to mosaic aneuploidy where the haplotypes may not exist as integer copies due to heterogeneity in somy of cells. Therefore, we extended POPSICLE, which we previously built for haploid genomes to accommodate aneuploid genomes (Shaik et al. 2018). We did this by leveraging allele frequencies to capture non-integer haplotype assignments to detect shared ancestries across populations. We have utilized 50 publicly available WGS strains corresponding to 23 distinct species of *Leishmania* to study several aspects such as genomic divergence, aneuploidies, single nucleotide polymorphisms, genome-wide zygosity profiles and correlations between somies and allele frequency distributions. The validity of POPSICLE in revealing admixtures in aneuploid genomes to understand the sexual strategies in *Leishmania* are discussed further. The strategies employed in this work can be directly applied to other mosaic aneuploid genomes with Autism, Schizophrenia, Auto-immune diseases, Alzheimer’s, Cancers and *Fungi*.

## Results

### Leishmania are aneuploid in all the species and strains studied

*Leishmania* exhibit a high degree of synteny (Supplementary Figure 1) and have previously been shown to contain only a few species-specific genes (Ghedin et al. 2004). Phenotypic variety may therefore be due to aneuploidy that is now known to be prevalent in *Leishmania* (Rogers et al. 2011). We sought to obtain a snapshot of aneuploidy in *Leishmania* to verify whether aneuploidies correlated with species/strain assignments. We aligned the short reads from Illumina sequences to the closest available reference sequence by employing BWA and using default parameters (Supplementary File1) (Li and Durbin 2009). We counted the reads that aligned to each chromosome and translated them into somies. The chromosomes were generally disomic, but deviations were observed in a handful of chromosomes (Figure 1). Chromosome 31 (chromosome 30 in *L. mexicana*) consistently showed an elevated somy of 4 or more. Likewise, chromosome 23 also routinely showed somies greater than 2 in many strains. Aneuploidy on the other chromosomes appeared to be random and showed no correlation with either species or strain assignments. *Leishmania* are also known to have non-integer somy values because of mosaic aneuploidy as the cells in a clonal population can exist in a variety of somy states and the reckoned somies are an average of all the somies present (Sterkers et al. 2012). Despite such genome plasticity, we observed most chromosomes were disomic on average.

**Figure 1.**
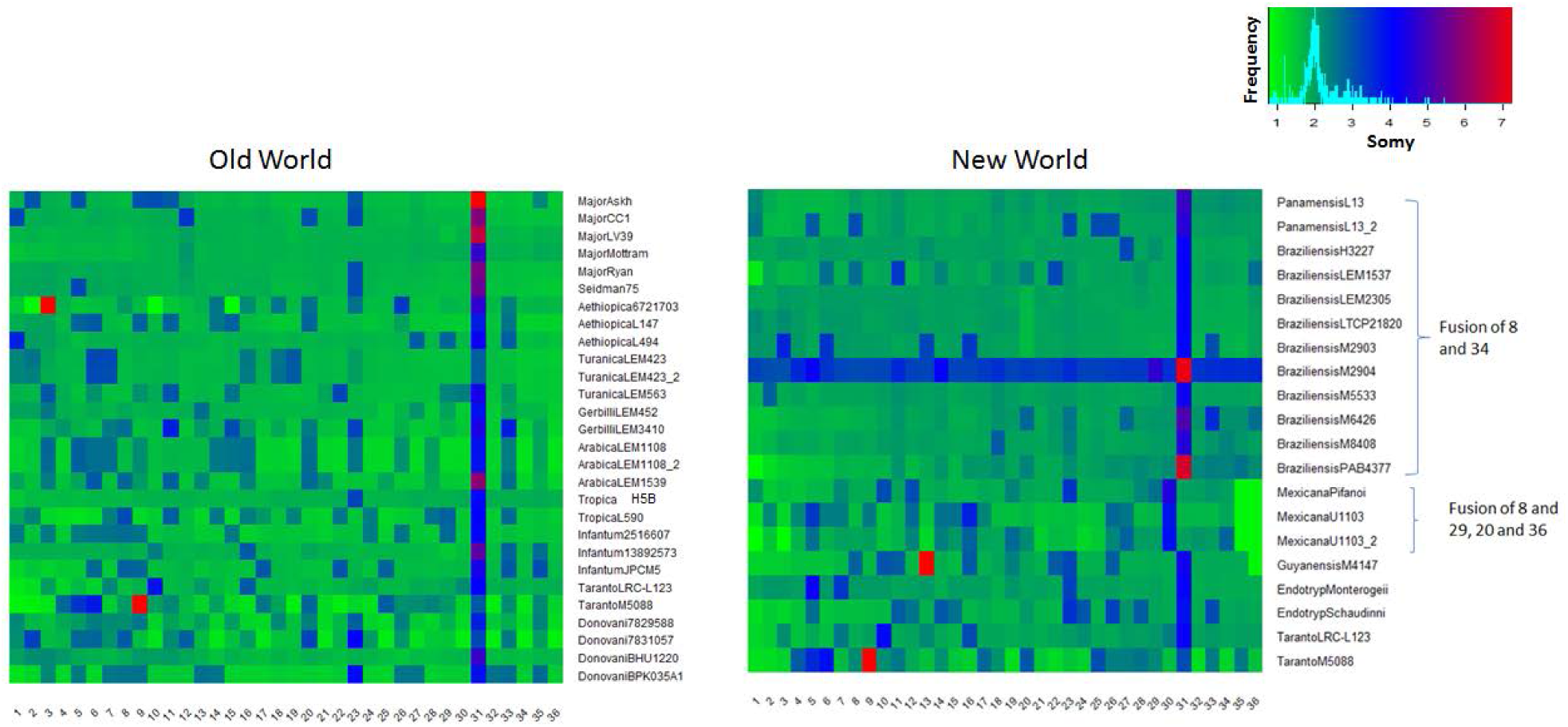
Aneuploidies in *Leishmania* determined by translating read mappings to somies. All strains are considered diploid except *M2904* which is known to be triploid. The somies are depicted using the indicated spectrum of colors. Most chromosomes are disomic on average, irrespective of species or strain (green). No correlation between aneuploidy and species assignments is observed, rather, aneuploidy seemed to be random. Only 34 chromosomes exist in *L. mexicana* and 35 in *L. guyanensis* in comparison to 36 in most of the strains because of various chromosome fusions.

**Supplementary Figure 1.**
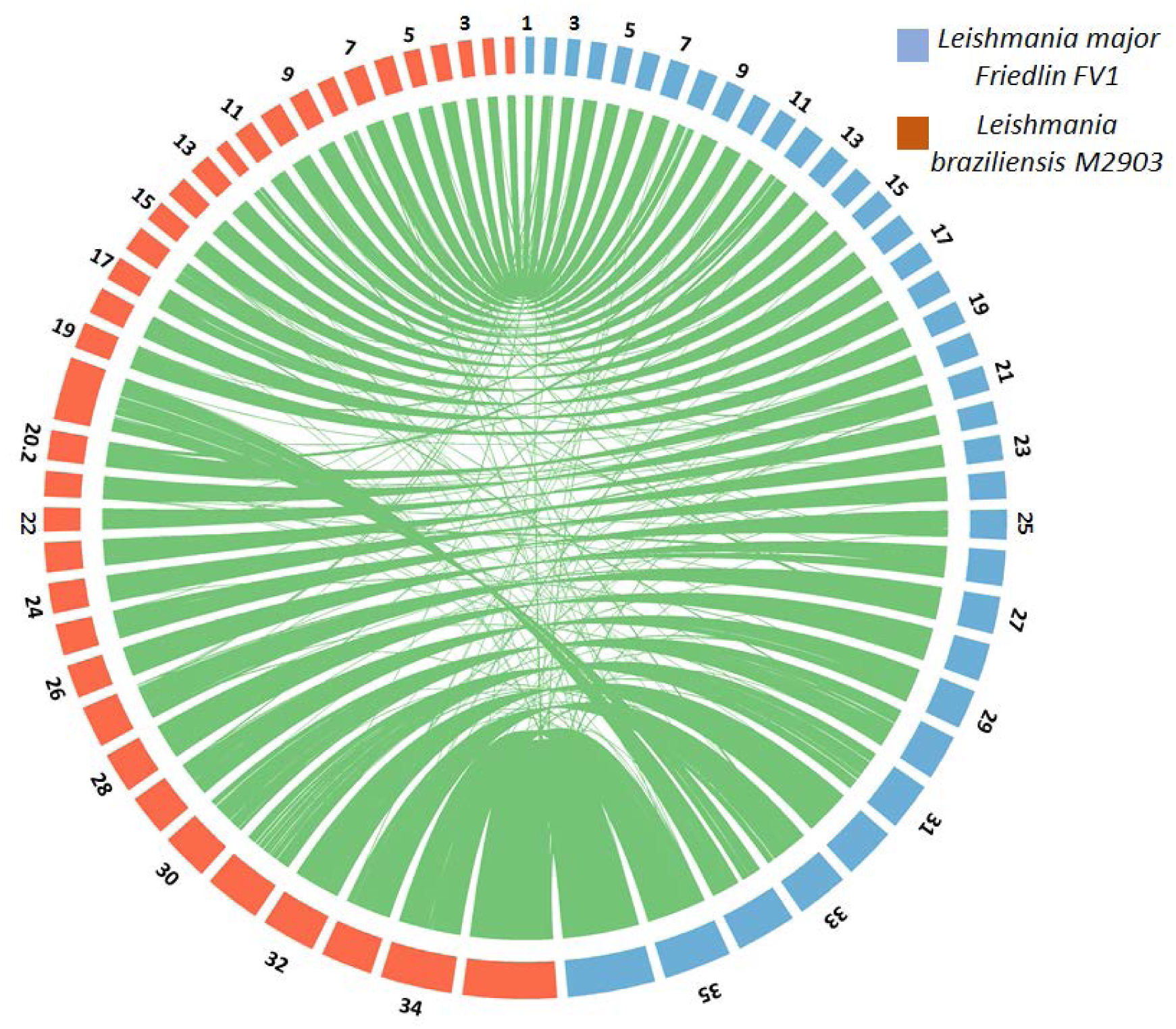
Synteny map between *L. major Friedlin FV1* strain from old-world belonging to Leishmania subgenus and *L. braziliensis M2903* strain from new world belonging to Viannia subgenus shows both genomes are highly syntenic

### Leishmania are diverse and form clades that agree with species assignments

*Leishmania* are extremely diverse and we sought to verify whether this diversity is reflected in the genomic profiles and if these profiles correlated with the species assignments. We did this by mapping the high-resolution NGS data against the well annotated *Leishmania major FV1* genome (Tritrypdb LmjF Version 33-LmjF33). We obtained alignment percentages with low alignment numbers (average 27%) for *Viannia* subgenus and high alignment numbers (average 95%) for *L. major* species (Supplementary File2) based on the divergence of the strains from the reference. We found single nucleotide polymorphisms (SNPs) using the samtools utility (Li et al. 2009) (described in Methods) against the LmjF33 and found anywhere between 714 and 2439280 SNPs in individual strains (Supplementary File2). As we were interested in broader variations across these strains, we consolidated these SNPs, removed the private SNPs (SNPs specific to individual strains), filtered out markers which had missing data in 30% or more strains and utilized 6364002 remaining markers to assess divergence in *Leishmania* using conventional dimensionality reduction methods such as principal component analysis (PCA) and hierarchical clustering. PCA analysis revealed the diversity between strains at a subgenus level (Figure 2A). The strains from the *Enriettii* complex were the most distinct followed by the strains from the *Paraleishmania* subgenus. *Endotrypanum* and *Sauroleishmania* were the most closely related to the *Leishmania* subgenus. The *L. mexicana* complex from the new world distinguished itself from the old-world species within the *Leishmania* subgenus. The new world strains from the *Viannia* subgenus formed a separate, distinct cluster. The phylogenetic analysis using hierarchical clustering framework with complete linkage revealed that strains belonging to the same species and species belonging to a sub-genus generally grouped together, although some exceptions were seen (Figure 2B). The species *L. tropica* and *L. aethiopica* formed a subgroup that suggested close association between them, a finding consistent with what previously been reported in a comparative analysis performed using microsatellite markers (Krayter et al. 2015). Similarly, the *L. donovani* complex containing *L. infantum* and *L. donovani* were assigned to a single clade, signifying their genomic similarity as reported previously (Lukes et al. 2007). The strains belonging to species *L. major, L. turanica, L. gerbilli* and *L. arabica* formed distinct subgroups as expected. The other strains from the *Viannia* subgenus were assigned to a single clade although subclades differentiated them by species. The only exceptions were two strains, *L. infantum* (ERR984221) which clustered with the *L. mexicana* complex and *L. pifanoi* Ltrod (SRR1662200) from the *Viannia* subgenus which clustered with *L. mexicana*. The *Endotrypanum, Paraleishmania* and *Enrietti* complexes are subgenera distantly related to the majority of *Leishmania* and these formed individual clades.

**Figure 2.**
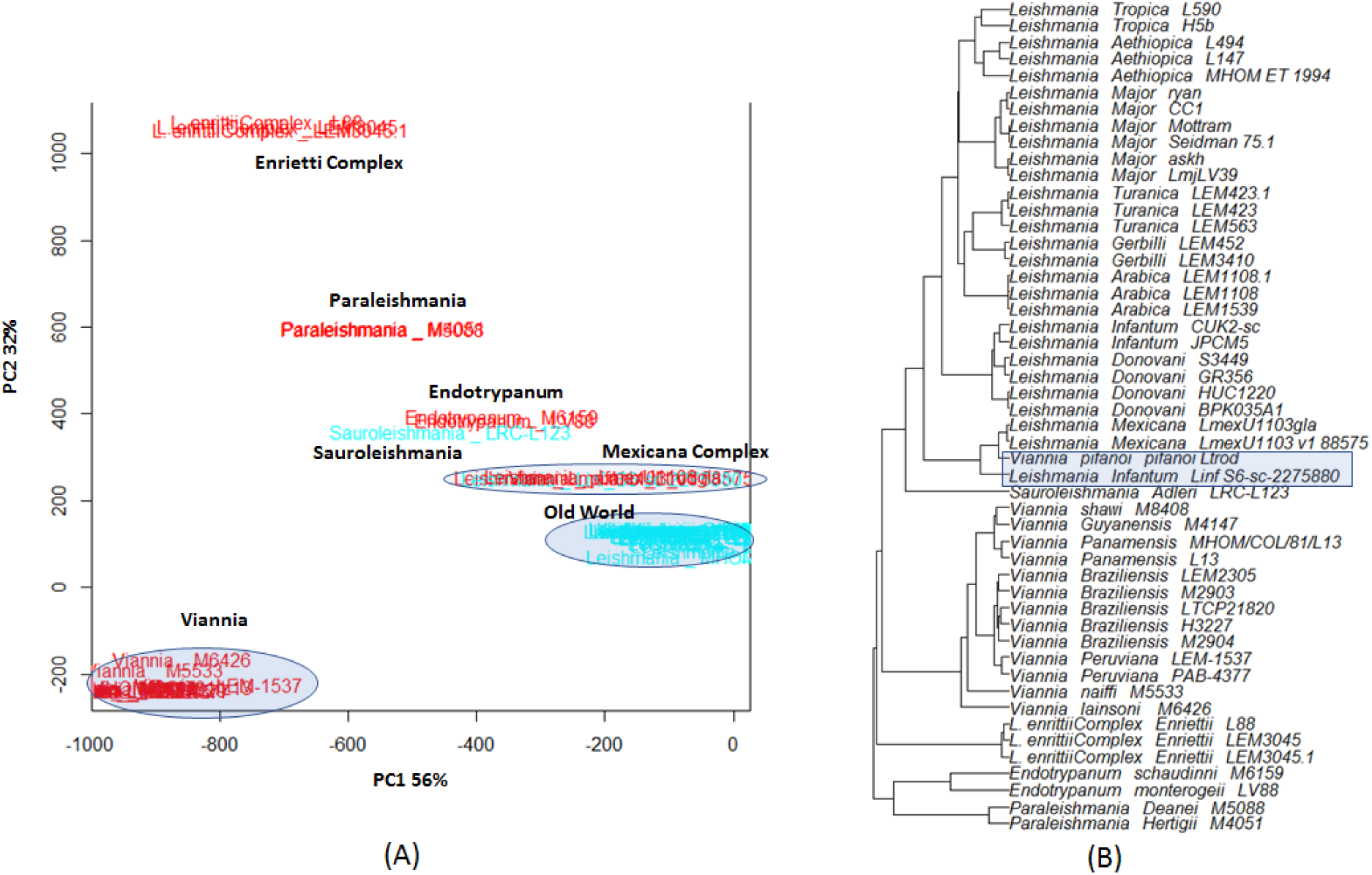
The phylogenetic analysis of various *Leishmania* species performed by leveraging read alignments to a common reference genome (LmjF33). A) Principal component analysis (PCA) of the data reveals a broad categorization of the strains at a subgenus level. New world (red) and old world (blue) species are indicated. Samples occupy different locations within the 2D space and are labelled using black text. B) Hierarchical clustering of the strains shows clustering by species. The clustering of most samples was as expected except for the highlighted strains. These strains can also be seen clustering with the *L. mexicana* complex in the PCA plot.

### Leishmania exhibit extreme hetero or homozygosity

The WGS reads of available *Leishmania* strains were aligned to the closest available reference sequence using BWA (Li et al. 2009) and obtained between 70% and 99% alignment rates (Supplementary File1). We enumerated the proportion of homozygous and heterozygous SNPs in *Leishmania* and found that 20 strains (42%) contained a high proportion of homozygous SNPs, 22 strains (46%) mostly contained heterozygous SNPs and 5 strains (12%), mainly belonging to *L. braziliensis* contained equal percentages of homozygous and heterozygous SNPs (Figure 3 and Supplementary File1). We were intrigued by the fact that most *Leishmania* showed extreme bias in zygosity. We investigated whether this bias was limited to a certain subgenus, geography or clinical phenotype and found no strong correlation with any of these factors using a linear regression model. These patterns of extreme zygosity were observed in both new and old-world strains. To visualize the location of these SNPs genome-wide, we enumerated the percentage of homozygous and heterozygous SNPs in sliding blocks of 5kb across the genome. We found that individual strains predominantly contained long runs of heterozygous or homozygous blocks (Figure 4). We observed negligible blocks of alternate zygosity in those strains with long runs of homozygosity. In contrast, strains with long runs of heterozygosity contained some blocks of alternate zygosity, sometimes spanning over entire chromosomes. These patterns of zygosity may provide a snapshot of introgressions in *Leishmania*; the homozygous strains may have hybridized to produce the heterozygous strains and the strains with balanced zygosity may have been produced by the mating between the heterozygous and the homozygous strains (Supplementary Figure 4). Although a few exceptions are seen, in general, *L. aethiopica, L. arabica, L. enriettii, L. gerbilli, L. guyanensis, L. mexicana, L. monterogeii* and *L. turanica* were heterozygous. In contrast, *L. adleri, L. deanei, L. donovani, L. infantum, L. liansoni, L. major, L. naiffi, L. panamensis, L. peruviana, L. pifanoi, L. schaudini* and *L. shawi* were predominantly homozygous.

**Figure 3.**
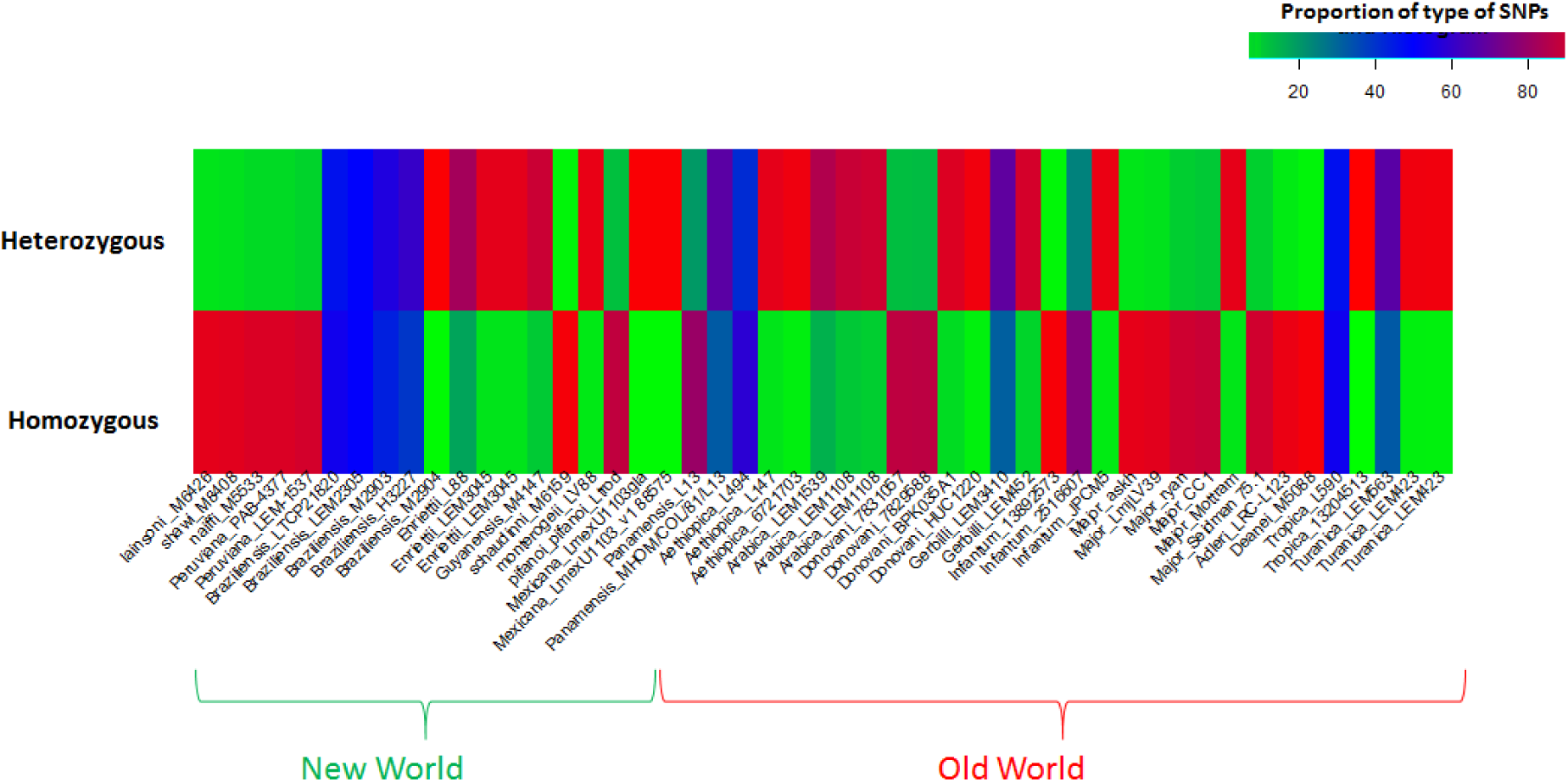
Proportion of homozygous and heterozygous SNPs in *Leishmania* show dichotomy in zygosity profiles. Proportions are represented using the indicated spectrum. Most strains either have high proportion of homozygous SNPs (red) and consequently low proportion of heterozygous SNPs (green) or vice versa. A few strains, such as from *L. braziliensis* contain 50% homozygous and 50% heterozygous SNPs (blue).

**Figure 4.**
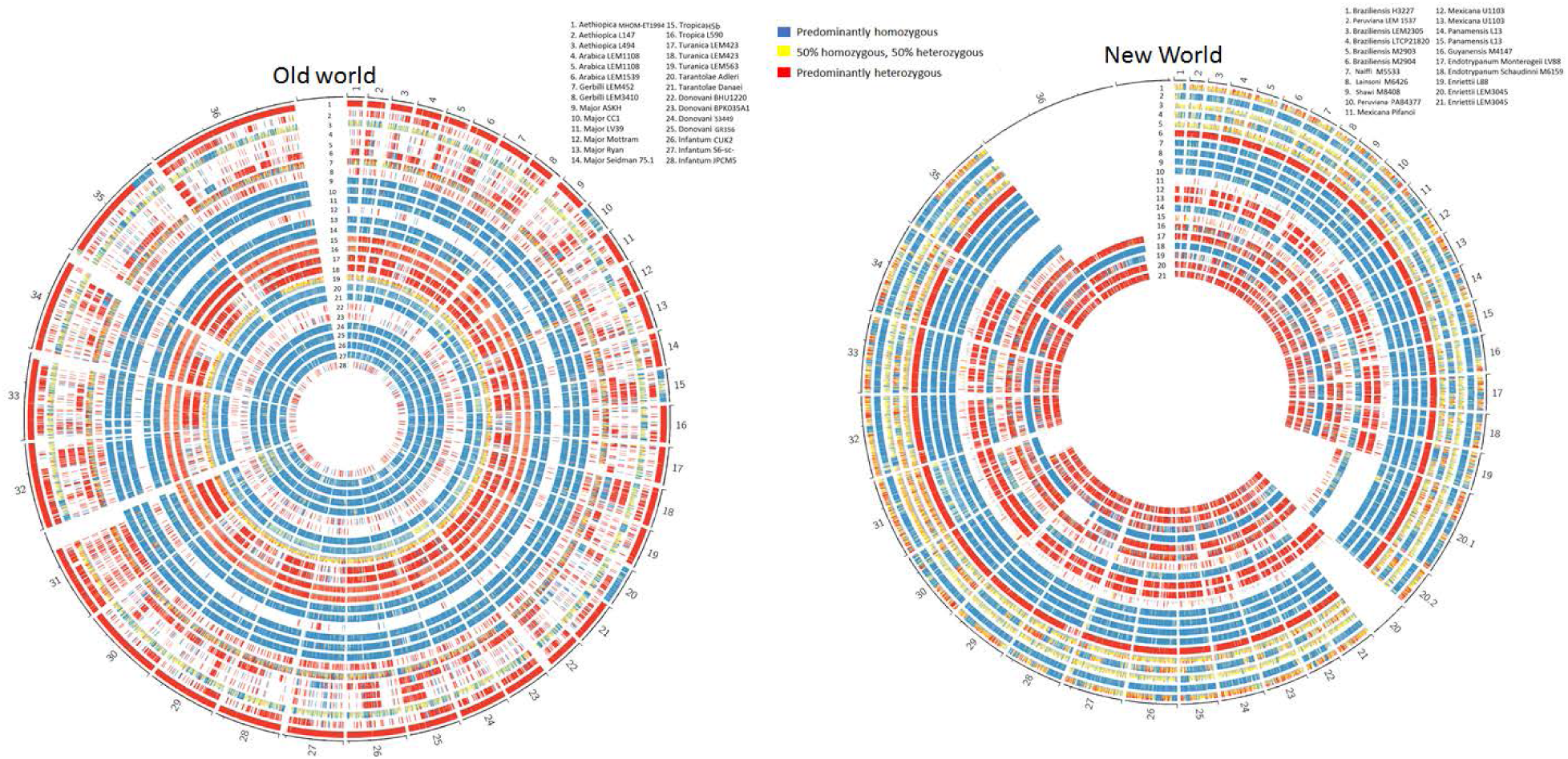
Genome-wide distribution of heterozygous and heterozygous SNPs in 28 old-world strains and 21 new-world strains of *Leishmania* predominantly contain long runs of heterozygosity or homozygosity. The zygosities were enumerated in blocks of 5kb and colored blue if the percentage of homozygous SNPs were greater than 80%, as red if the percentage of heterozygous SNPs was greater than 80% and as yellow if otherwise. Only a few strains contain long runs of balanced zygosity. The homozygous strains contain negligible blocks of alternate zygosity whereas the heterozygous strains contain blocks of homozygosity or balanced zygosity across the genome.

### The allele frequency distributions of the heterozygous SNPs in Leishmania agree with somies in some strains but do not agree with somies in some others

The distribution of heterozygous allele frequencies must reflect the somies of the chromosomes. In a disomic chromosome, a heterozygous SNP represents an alternate allele on one of the two haplotypes. The regions containing heterozygous alleles when sequenced generate 50% of the reads that are consistent with the reference allele representing one haplotype and the other 50% that match the alternate allele on the other haplotype. Although minor differences in percentages that can be attributed to the factors such as bioinformatics error are expected, in general, these percentages are close to 50%. The distribution of the heterozygous allele frequencies therefore peaks around the value of 0.5. Similarly, on trisomic chromosomes, the allele frequencies peak around 0.33 and 0.66 and around 0.25, 0.75 and 0.5 on tetrasomic chromosomes. The allele frequency distributions therefore provide important clues regarding the somy of a chromosome. We categorized chromosomes as disomic if the somy was between 1.75 and 2.25 and as trisomic if the somy was between 2.75 and 3.25 and drew allele frequency distributions. We noticed that a few strains, as expected, had strong correlations between the somies and the allele frequency distributions while some did not. For example, the triploid *L. aethiopica* strain MHOM ET 1994 from the old-world show peak at 0.5 for disomic chromosomes and at 0.33 and 0.66 for trisomic chromosomes as expected (Figure 5A). The tetrasomic chromosomes either show peaks at 0.25, 0.5 and 0.75 indicating more than 2 haplotypes or at 0.5 indicating 2 haplotypes. The hexasomic chromosome shows peaks at 0.17 and 0.83 indicating presence of only two haplotypes. Similarly, in the new-world, *L. braziliensis* M2904 shows peaks near 0.33 and 0.66 for trisomic and three peaks near 0.25, 0.5 and 0.75 for tetrasomic chromosomes indicating more than 2 haplotypes. The hexasomic chromosome 31 shows a complex pattern with peaks near 0.166, 0.33, 0.5, 0.66 and 0.88, also indicating more than 2 haplotypes. However, there were some strains such as *L. major* ASKH that showed noisy allele frequency distributions that don’t match the somy (Figure 5c). These disagreements between somies and allele frequencies have been seen predominantly in homozygous strains with low SNP counts.

**Figure 5.**
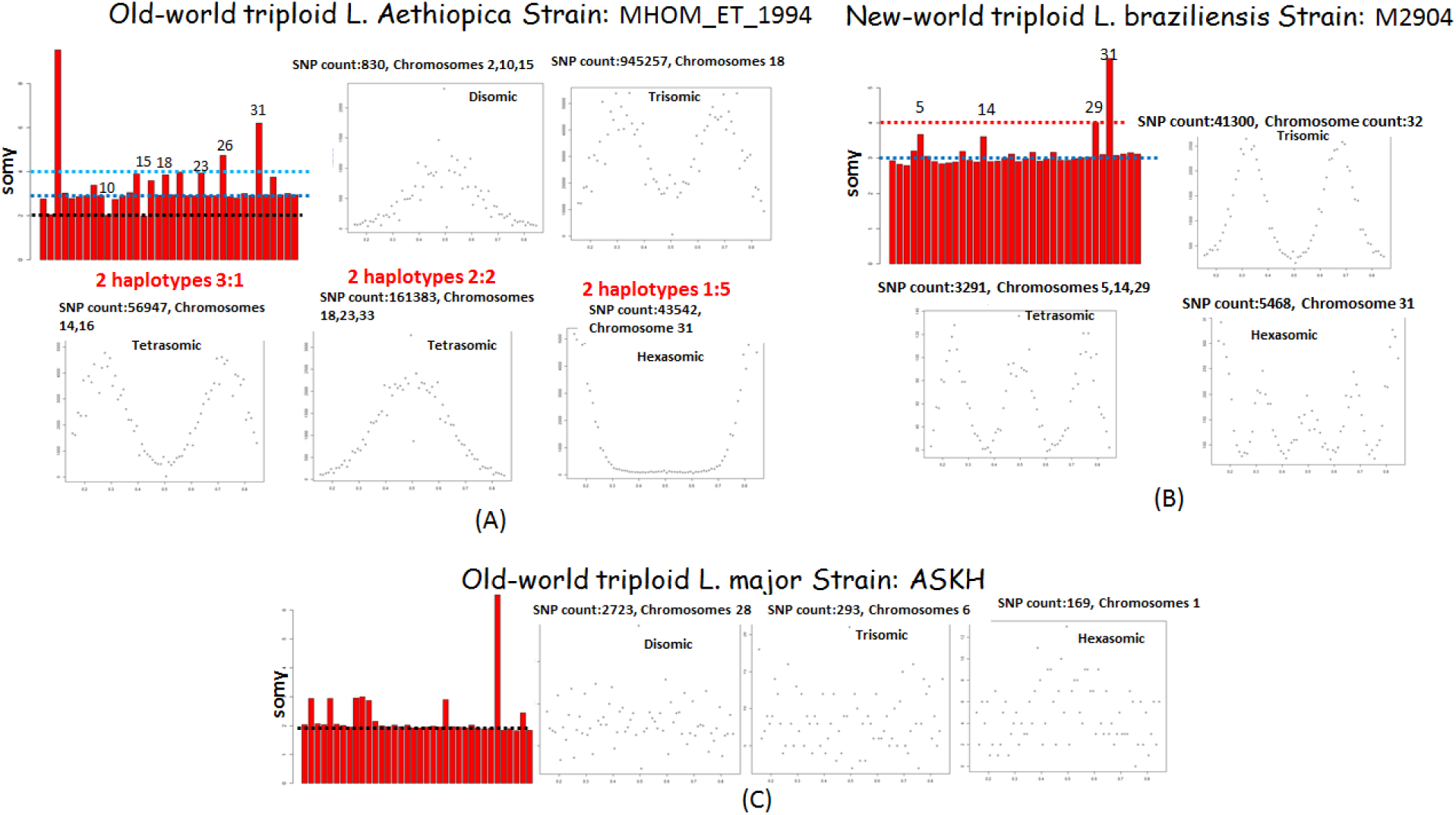
Concordance between the somies and allele frequency distributions. A) The *L. aethiopica* strain MHOM_ET_1994 from old-world and *L. braziliensis* strain M2904 from new-world are triploid based on allele frequency distributions. The somy distribution of MHOM_ET_1994 reveals 18 trisomic, 3 disomic, 5 tetrasomic and one hexasomic chromosomes. The chromosome numbers for aneuploid chromosomes are indicated on top of the bars. B) Similarly, the somy profile of M2904 shows 32 trisomic, 3 tetrasomic and one hexasomic chromosomes. The allele frequency distributions of the chromosomes from these two strains match the respective somy profiles. The strain MHOM_ET_1994 contains only 2 haplotypes whereas the strain M2904 contains more than 2 haplotypes as evident from the allele frequency distributions of tetra and hexasomic chromosomes. C) Some of the strains however do not show correlation between allele frequency distributions and somies as shown for *L. major ASKH*. The allele frequency distributions in this strain have noisy patterns irrespective of the somy.

### Population Structure of Leishmania

In order to reveal the extent of shared ancestries among the various species of *Leishmania*, particularly among closely related species, we employed POPSICLE for a more in-depth analysis using the 6364002 markers, which we had previously employed to draw phylogenies, as our input data. POPSICLE assigned the samples to 22 clades which were mostly in agreement with the species assignments with a few exceptions that were also observed in the phylogenetic analysis (Figure 6). To verify if this misclassification was due to the hybrid nature of the strains, mixed infections or due to mislabeling of samples, we tested the *L. aethiopica* strain MHOM ET1994 which had clustered with *L. tropica*. Chromosomes 10, 15 and 20 of MHOM ET1994 were similar to *L. aethiopica* but chromosome 35 was partially heterozygous and partially homozygous. POPSICLE indicated that the heterozygous segment was inherited from *L.tropica* whereas the homozygous segment was from *L. aethiopica* (Figure 6). This suggested that although classified as *L. aethiopica*, MHOM ET1994 was a hybrid. We investigated this further using homozygous SNP differences between *L. aethiopica* and *L. tropica* genomes (253244 markers), an analysis that revealed that MHOM ET1994 inherited alleles from both L. *tropica* and *L. aethiopica* as expected from an outcross. Although mostly heterozygous, the strain contains a few segments with loss of heterozygosity (LOH) that looked like *L. aethiopica* (supplementary Figure 2A). Hybrids in *Leishmania* sometimes contain regions with LOH that are not confined to backcrosses as were previously observed in intra and inter-species experimental F1 hybrids (Inbar et al. 2013). The frequency of the observed LOH in MHOM ET1994 was high (11%) in comparison to what was previously observed in F1 experimental hybrids (<1%) suggesting that MHOM ET1994 is not a F1 hybrid but instead a product of multiple cycles of inter-species outcrossing.

**Figure 6.**
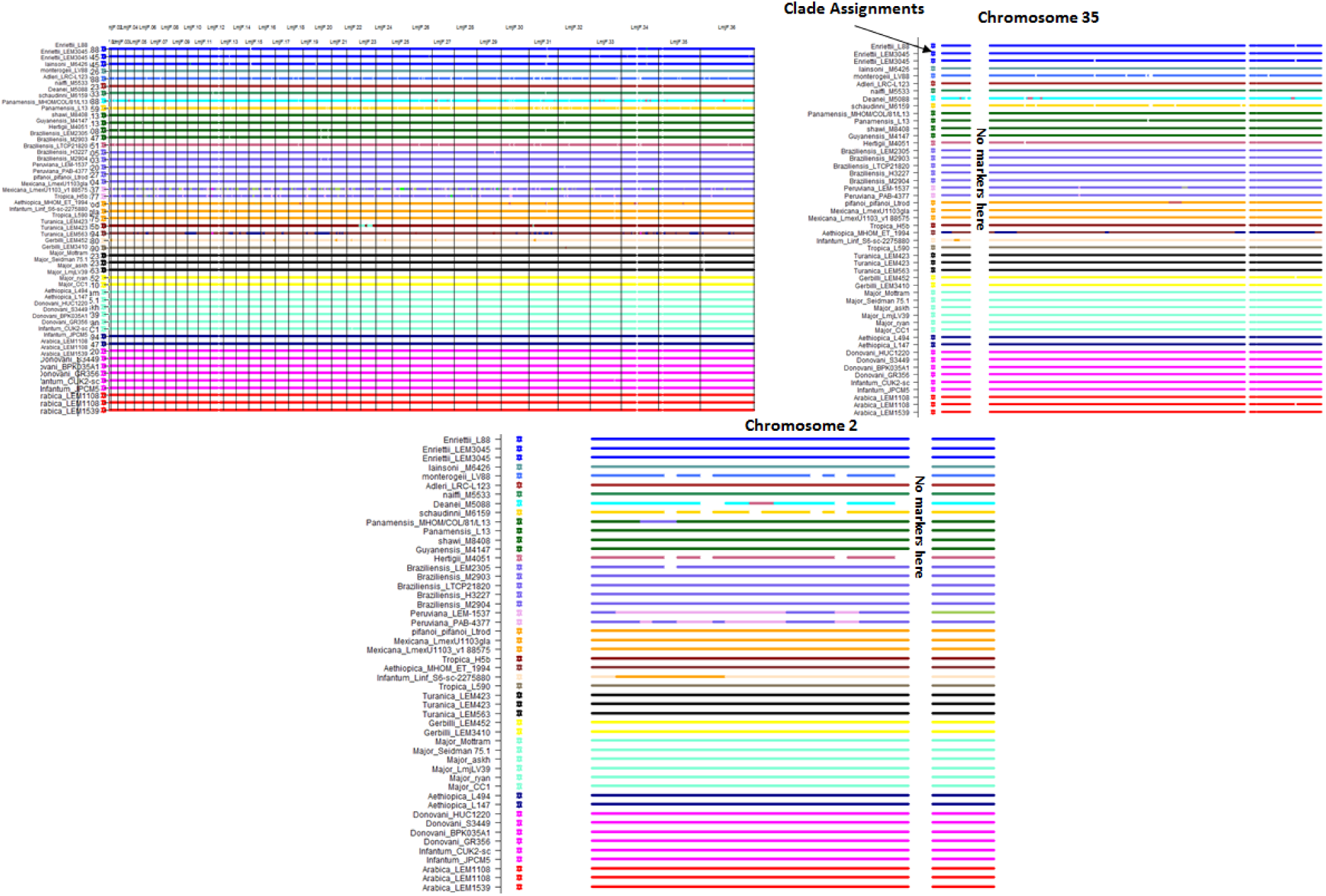
Population Structure of *Leishmania* captured using POPSICLE. The clade assignments of each strain are indicated by colored stars at the left of profiles. The strains in general contain the color of the clades assigned to them except where they share ancestries with other strains. Population structure reveals associations at a species level. Most strains were true to their clade assignments with only a few blocks of shared ancestries. Exception to this rule were strains such as *L. infantum* S6-SC-2275880 that contain blocks of shared ancestry with *L. mexicana*. Shared ancestries between closely related species (e.g. *L. peruviana* and *L. braziliensis)* can be observed at a chromosome resolution as shown for chromosomes 35 and 2. Chromosome 35 in MHOM_ET_1994 is partially heterozygous and partially homozygous (Figure 4). POPSICLE indicated that the heterozygous segment of the genome is inherited from *L. tropica* whereas the homozygous segment is inherited from *L. aethiopica* suggesting its hybrid nature.

We also investigated the homozygous *L. infantum* strain S6-sc-2275880, which did not phylogenetically cluster with the other *L. infantum* strains, instead grouping with *L. mexicana*. POPSICLE indicated that it has a peculiar profile and contained many intermittent blocks of shared ancestry with *L. mexicana*, for example on Chromosome 2 (Figure 6). This strain exhibited long runs of homozygosity indicating that it is likely not a product of recent matings as we would expect it to be largely heterozygous strain similar to MHOM ET1994. To confirm this, we examined the 155763 homozygous SNP differences between *L. mexicana* and *L. infantum* genomes. This analysis confirmed that S6-sc-2275880 is predominantly *L. mexicana* although a few traces of *L. infantum* are seen (supplementary Figure 2B). These results are most likely due to a coinfection but extreme inbreeding involving *L. mexicana* and *L. infantum* cannot be ruled out as a possibility. Although phylogenetic analysis indicated that the two strains MHOM ET1994 and S6-sc-2275880 did not cluster with the species they were expected to, the additional insights gained using POPSICLE provides a level of detail without which such categorizations would have been difficult to explain.

**Supplementary Figure 2.**
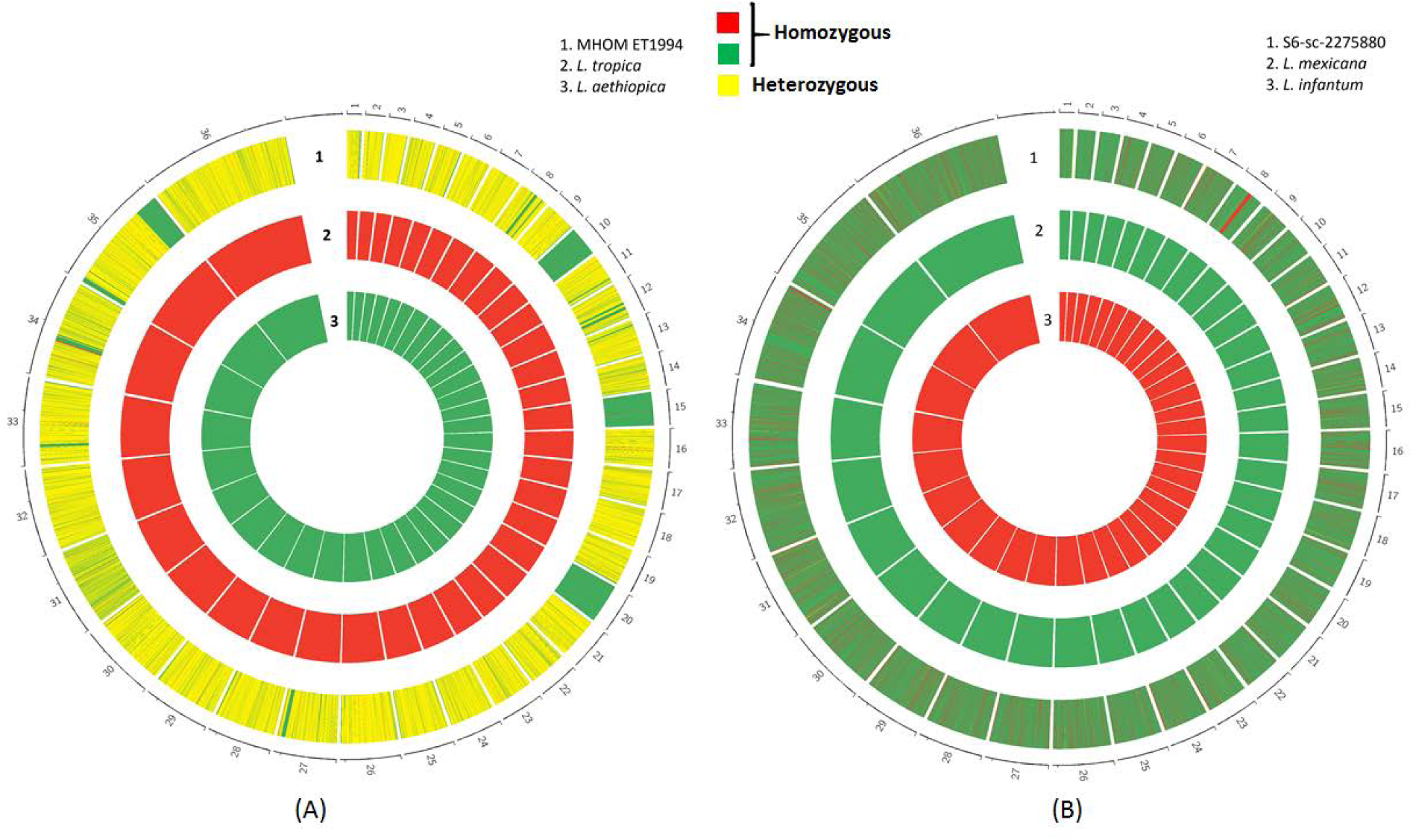
Allele compositions of the strains misclassified using hierarchical clustering. (A) Markers that are homozygous and different between *L. tropica* and *L. aethiopica* are considered for this analysis. Homozygous alleles in *L. tropica* are shown in red, those in *L. aethiopica* are shown in green. MHOM_ET_1994 is mostly heterozygous (shown in yellow) and inherited alleles from both *L. tropica* and *L. aethiopica* suggesting its hybrid nature. A few homozygous blocks of genotypes matching *L. aethiopica* are also seen. B) A similar strategy was followed to analyze the strain S6-sc-2275880 which is classified as *L. infantum. L. mexicana* was employed as the other putative parent as S6-sc-2275880 clustered with *L. mexicana* (Figure 2). Homozygous alleles in *L. infantum* are colored red and those in *L. mexicana* are colored green. The profile of S6-sc-2275880 indicates that it mostly contains alleles from *L. mexicana* and a few blocks of *L. infantum* thereby justifying its clustering with *L. Mexicana* and not with *L. infantum.*

### Zygosity and population structure of L. Tropica reveals genotypes that correlate with geography

The two *L. tropica* strains (L590 and H5b) that we initially analyzed contained a substantial number of heterozygous SNPs distributed across the genome. We therefore probed a larger collection of 19 strains to determine whether all *L. tropica* are heterozygous. We aligned the short reads corresponding to these strains to *L. tropica* genome (TrytrypDB L590 V. 33- Ltr33) and achieved between 71% and 90% alignment rates and between 104,000-284,000 SNPs (Supplementary File3) which were employed as markers for our analyses.

### A variety in zygosity is observed in L. tropica and it correlates with the geographic origin

We enumerated heterozygous and homozygous SNPs in these 19 strains and found that unlike the two strains that were initially studied, 8 (42%) were predominantly homozygous, 10 (52%) were heterozygous and 1 (5%) contained equal numbers of heterozygous and homozygous SNPs (Supplementary File3). We computed the genome-wide distribution of these SNPs by counting the percentage of heterozygous and homozygous SNPs in blocks of 5kb across the genome for all strains (Figure 7A). The strains with predominantly homozygous SNPs showed long runs of homozygosity with negligible blocks of alternate zygosity. The strains with largely heterozygous SNPs in contrast showed long runs of heterozygosity and against this background were the blocks of homozygosity, sometimes spanning over the entire chromosome. For example, in the Ackerman strain of *L. tropica*, against a heterozygous background, chromosomes 8, 9, 10, and 34 were entirely homozygous and chromosomes 1, 11, 22, 29 and 36 contained large blocks of homozygosity. A strong correlation between zygosity and geographical origin of the strains was observed. Those strains with extreme heterozygosity were spread over a wide geographical area encompassing India, Afghanistan, Iraq and Syria (Figure 7B). In contrast, strains with extreme homozygosity were restricted mainly to a small geographical area that included Palestine, Israel and Jordan. Strains from Saudi Arabia were mainly heterozygous but contained blocks of homozygosity, whereas the strain from Tunisia were mainly homozygous with a few blocks of balanced zygosity.

**Figure 7.**
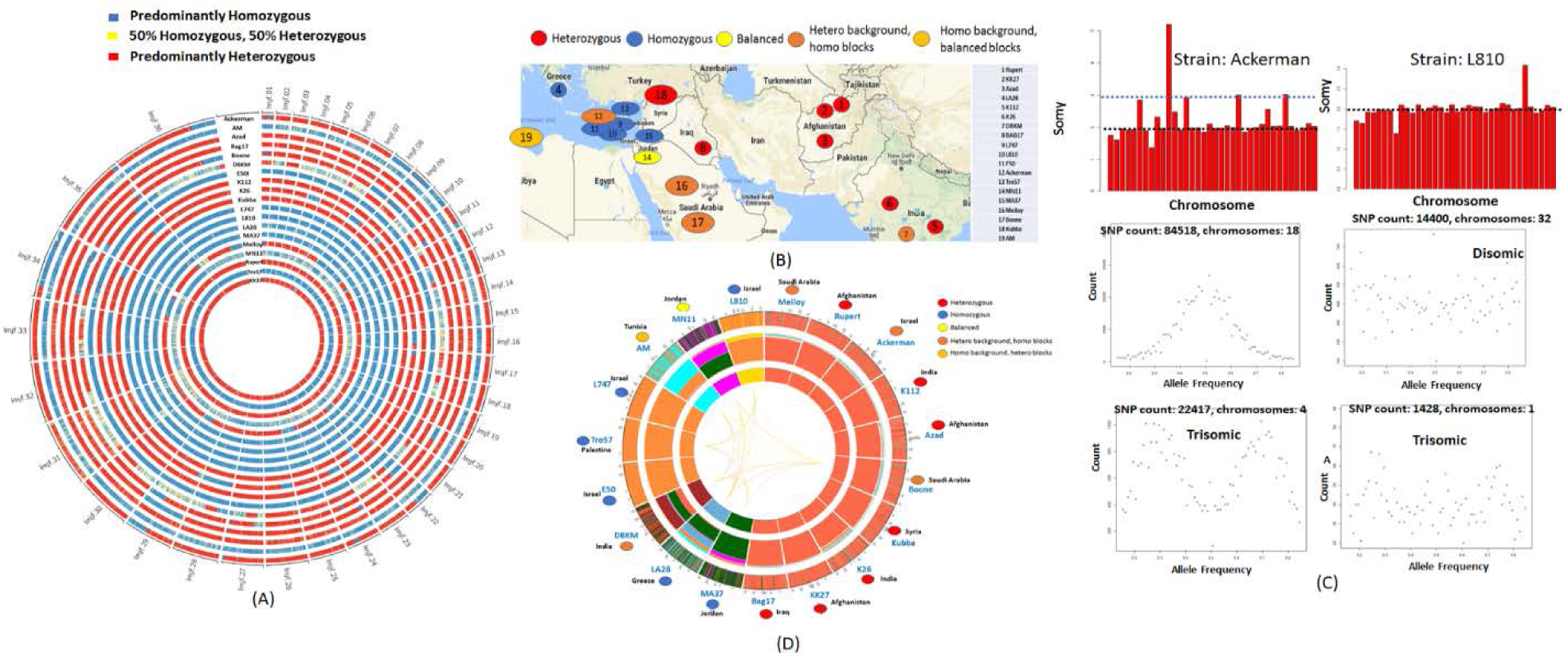
Diversity in *L. tropica*. (A) The proportion of heterozygous and homozygous SNPs in blocks of 5kb are calculated and colored as indicated in the legend. The zygosity profile of *L. tropica* indicates long runs of heterozygous or homozygous blocks. B) The zygosity profile correlates with the geography. The strains from India, Afghanistan, Syria and Iraq are mainly heterozygous. The strains from Israel, Palestine and Jordan are mainly homozygous. C) The allele frequency profiles do not agree with somy in all the strains. The agreement is strong in strains with extreme heterozygosity and is minimal in strains with extreme homozygosity coupled with low SNP count. *Ackerman* which contains 89% heterozygous SNPs is diploid as suggested by its ploidy profile. The allele frequency profile of 18 disomic chromosomes shows a peak near 0.5 and those of 4 trisomic chromosomes shows peaks near 0.33 and 0.66 as expected. The strain L810 which contains 97% homozygous SNPs is also diploid but the allele frequency profiles irrespective of somy show noisy patterns. D) Overview of the population structure in *L. tropica*. The inner concentric circle depicts the clade assignments and the extent of shared ancestries between the clades are indicated by the links that connect them. The middle concentric circle depicts the global ancestral profiles that show proportion of shared ancestries of a strain with other clades. The outer concentric circle depicts the genome-wide ancestral profiles of the strains.

### The allele frequency distributions sometimes do not match the somy in L. tropica

We verified whether the allele frequency distributions agree with the chromosomal somies in *L. tropica* using our previously described criteria and studied their distribution. While 13 strains (68%) of *L. tropica* showed correlation between the somy and allele frequency distributions, 6 strains (32%) did not. The Ackerman strain for example shows a peak near 0.5 for disomic chromosomes and two peaks near 0.33 and 0.66 for trisomic chromosomes respectively as expected (Figure 7C). In contrast, the *L810* strain showed noisy patterns irrespective of the somy compositions. We determined that the strains that did not show the correlation between somies and allele frequency distributions were mainly homozygous and contained less than 6K heterozygous SNPs. The heterozygous SNPs in these strains are most likely a consequence of mosaic aneuploidy, which causes allelic differences between sub-populations within clonal population of cells, and not due to allelic differences among the homologous chromosomes.

### The ancestry profiles of L. tropica indicates four group of strains and reveals hybrids between two distinct homozygous strains

The SNPs from 19 strains against Ltr33 were consolidated to obtain 599032 total markers which were filtered for private SNPs and missing values, retaining 250634 markers. POPSICLE determined that there were 8 distinct clades to which all 19 strains were assigned. The samples which were mainly heterozygous were assigned to the red clade (Figure 7D-inner concentric circle). The homozygous strains from Israel and Palestine were assigned to the orange clade while the other homozygous strains were assigned to individual clades because of their unique profiles. The proportion of shared ancestries between the clades are illustrated using the links that connect them at the center. The global ancestral profiles indicate that minor variations were seen in the heterozygous strains assigned to the red clade. The homozygous strains although assigned to multiple clades, can be broadly classified into two major groups; The first group consisting of MA37, LA28 and MN11 contained shared ancestries as shown in green and the second group consisting of Tro57, E50, L747, AM and L810 contain shared ancestries as indicated in orange. The sample DBKM is unique consisting of both green and orange suggesting that it is possibly an outcross between groups 1 and 2. We verified whether the heterozygosity of the red clade was a consequence of mating between strains from groups 1 and 2. We selected 82234 homozygous SNPs from MA37 against Ltr33 and drew the allele compositions of the strains. The profiles that emerged indicated two major groups of homozygous strains like those observed in the ancestral profiles obtained using POPSICLE (Figure 8). These genotypic groups were associated with two different countries. The first group consisting of MA37, LA28 and MN11 mostly belonged to Jordan and the second group consisting of Tro57, L810, L747, E50 and AM mainly belonged to Palestine. The strains from Group 3 were the products of one cycle of mating between the strains from Jordan and Palestine whereas the strains from Group 4, were the products of multiple cycles of mating (Figure 8). Although the exact parents are unknown, one of the parental strains likely was of genotype such as from group1 and the other parent was of genotype similar to group 2.

**Figure 8.**
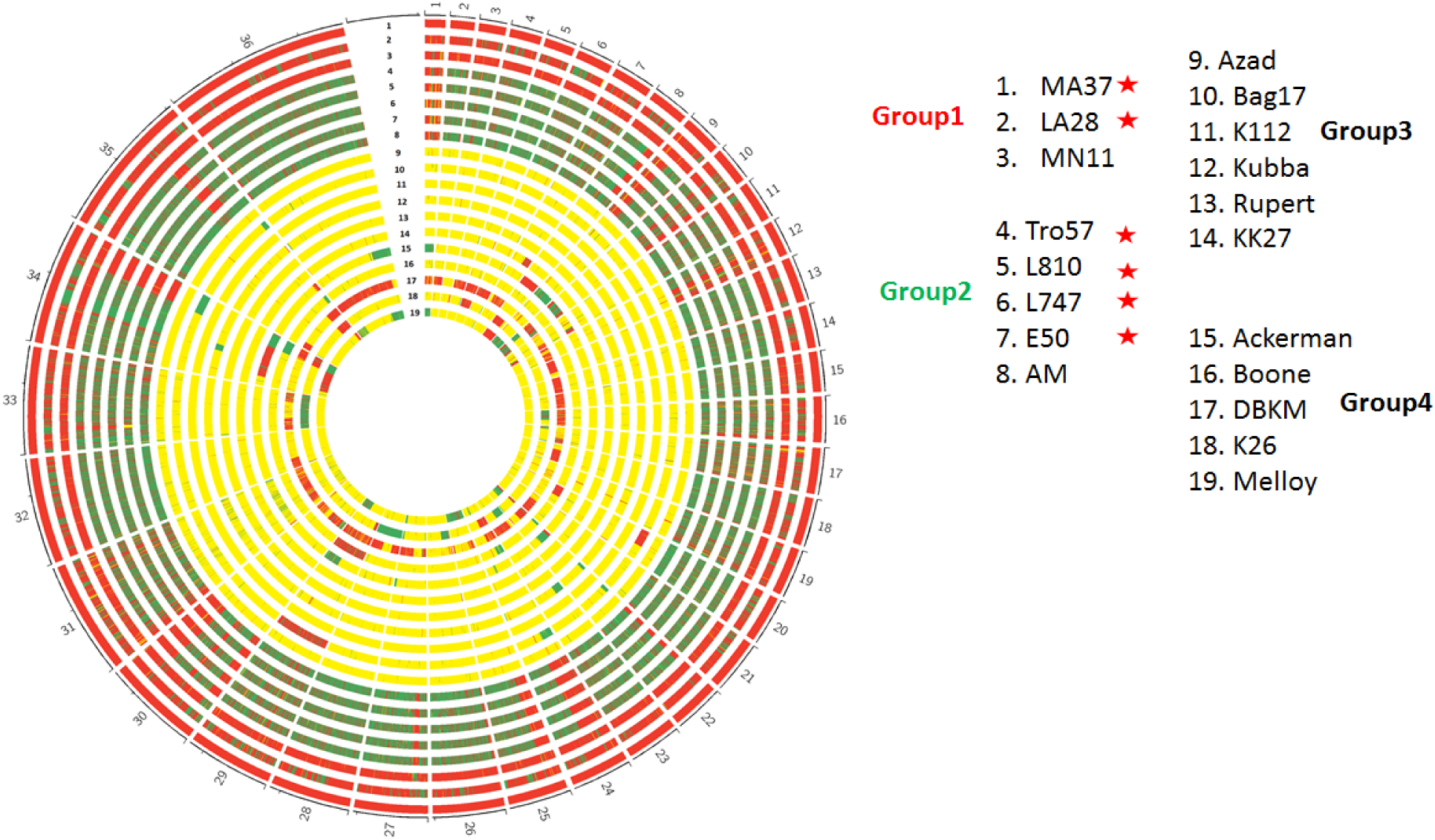
Allele compositions in *L. tropica* depicted using the markers where MA37 was homozygous. The alleles that match the strain MA37 were colored red, alternate homozygous alleles were marked green and heterozygous alleles consisting of both red and green alleles were colored yellow. Similar to what was suggested by POPSICLE, there were two major homozygous groups, group1 and group2. Group3 and group4 were the outcrosses involving groups 1 and 2. Strains from group 3 were most likely F1 hybrids between groups 1 and 2. Strains from group 4 were products of more than one cycle of mating. The strains in which the allele frequency distributions don’t match the somy are indicated by red stars.

The strains which showed no correlation between the allele frequencies and the corresponding somies (Figure 8, denoted by a red star) were from groups 1 and 2. These were mainly homozygous strains with less than 6000 heterozygous SNPs. On the other hand, the strains that were the products of mating classified as groups 3 and 4 contained abundant heterozygous SNPs because they were outcrosses involving two different genotypes from groups 1 and 2. This suggests that *L. tropica* emerged in Palestine and Jordan as two distinct homozygous strains. The genomes of the strains from group 1 were mainly unique to group 1 with some blocks of group 2 genotype and occasional blocks of heterozygosity are observed, with the inverse being observed for group 2. These patterns occurred likely due to extreme inbreeding involving the two major genotypes and their products of hybridization. Patterns such as these have previously been argued to be products of multiple hybridizations in *L. infantum* (Rogers et al. 2014b). As these strains migrated into nearby regions, samples from group 4 involving recent but multiple hybridizations were observed in Iraq and Syria. These eventually migrated to other parts of the world such as Afghanistan and India where F1 hybrids are observed. It has been shown that F1 hybrids in *Leishmania* contain a few regions of LOH that are not confined to backcrosses (Inbar et al. 2013). The only exception was *DBKM*, which showed blocks of homozygosity that mainly matched the group 1 genotype suggesting that it was a backcross between a heterozygous strain and a strain from group 1. *DBKM* was not a product of crossing involving two heterozygous strains because large proportions of group 1 and 2 genotypes were not observed as one would expect from crossing between two heterozygous strains. This suggests that the homozygous group 1 genotype must be present in India, however we did not see based it represented in the strains analyzed. Another possible explanation is that *DBKM* is a recent transmission to India. Almost all heterozygous strains in India and Afghanistan remained heterozygous and if they had mated, we would expect greater genotypic variety than that observed in the strains analyzed. This speaks to the possibility that these heterozygous strains are sterile to each other. Extrapolating this observation, the strains from group 4 were perhaps products of multiple hybridizations involving heterozygous and homozygous strains. This seems likely due to the geographic proximity of group 4 with homozygous strains of groups 1 and 2.

### Zygosity and population structure of L. major indicate products of extreme inbreeding

We initially studied five different strains of *L. major* and observed that they were mainly homozygous. We extended this analysis to 7 different strains of *L. major* (some strains were repeated) and subjected them to similar analyses we performed on *L. tropica*. We aligned the short reads to Lmj33 reference and achieved between 77% and 95% alignment rates and between 653 to 80,000 SNPs which were employed as markers for our analyses (Supplementary File5). We counted the number of heterozygous and homozygous SNPs in windows of 5KB across the genome and depicted them as SNP densities (Figure 9A). The strains *FV1, Mr. Friedlin* and *FA1* had fewer, mainly heterozygous SNPs as these strains were the closest to reference genome. The heterozygous SNPs were not distributed throughout the genome but were clustered in a few regions as seen for chromosomes 12 and 26. The remaining strains had homozygous SNPs distributed throughout the genome but a propensity to localize in regions was still observed. POPSICLE analysis assigned the *L. major* strains to four different clades correlating with their SNP profiles (Figure 9B). The strains FV1, Mr. Friedlin and FA1 had a fewer SNPs and were assigned to the blue clade, the strains LV39 and ASKH which contained more than 60,000 SNPs were assigned to the green clade and the strains CC1 and Ryan with approximately 32,000 and SD with approximately 45,000 SNPs were assigned to the red and brown clades respectively. The global ancestry profiles generated using POPSICLE suggest that the red and brown clades are mixed genotypes that involved the strains from blue clade. To determine the genotypes associated with these strains, we used *FV1* as the reference strain and analyzed the zygosity. The patterns that emerged indicated two major genotypes similar to what was suggested by POPSICLE (Figure 10A). The strains *FV1, Mr. Friedlin* and *FA1* were mainly red genotype, the strains *LV39* and *ASKH* were the alternate green genotype with some blocks of red genotype and the strains CC1, Ryan and SD were mainly red genotype with blocks of green genotype. The strains *SD, CC1* and *Ryan* contain similar patterns with common blocks indicating a common origin of divergence from the red genotype except for a few regions as evident from chromosomes 32 and 33. This indicates that they underwent subsequent changes to differentiate into different strains. We eliminated the private SNPs and redrew the allele compositions to verify if the differences among the strains were due to independent mutations (Figure 10B). The SD strain now looks similar to CC1 after removing private SNPs indicating they were different as a consequence of independent mutations acquired over time. We next verified if the heterozygosity in *L. major* was because of allelic differences between the homologous chromosomes as seen in some *L. tropica* strains or due to somy differences among the cells within a clonal population (Sterkers et al. 2014). We did this by comparing the allele frequency distributions against the somies and found that none of the strains had agreement between somies and allele frequency distributions irrespective of the heterozygous SNP counts (Figure 9C). This suggests that although some of the heterozygous SNPs may have occurred due to allelic differences among the homologous chromosomes, most heterozygous SNPs were a consequence of mosaic aneuploidy.

**Figure 9.**
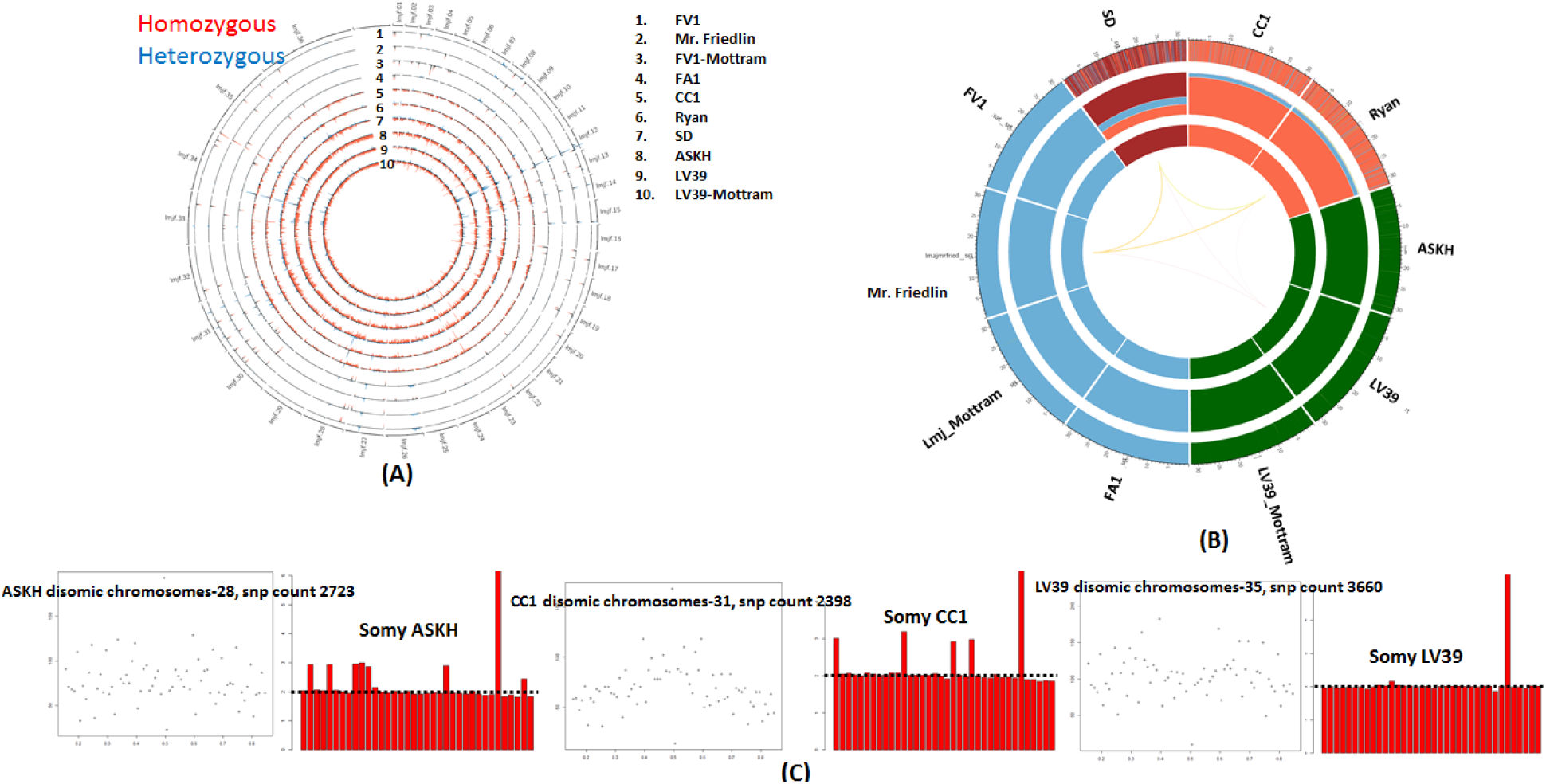
Population structure of *L. major* strains. A) The SNP density profile of *L. major* strains indicates low SNP counts, less heterozygosity and long runs of homozygosity as indicated by the red bars. The heterozygous SNPs were concentrated in certain segments of the genome (blue bars). B) Population structure obtained using POPSICLE indicates 4 clades. The global ancestry profiles (middle concentric circle) indicate that the strains SD, CC1 and Ryan have shared ancestries with the blue clade. The local ancestral profiles (outer concentric circle) show that these shared ancestries with the blue clade are not concentrated in a single region but are distributed throughout the genome. C) Since heterozygosity is very scarce in *L. major*, we elucidated if the observed heterozygosity is due to mosaic aneuploidy, we drew the allele frequency distributions of the heterozygous SNPs from the disomic chromosomes (majority of chromosomes are disomic) in strains that showed some heterozygosity. The allele frequency distributions of the disomic chromosomes do not show peaks at 0.5 as expected. Noisy frequency distribution patterns were similarly observed for chromosomes with other somies (data not shown).

**Figure 10.**
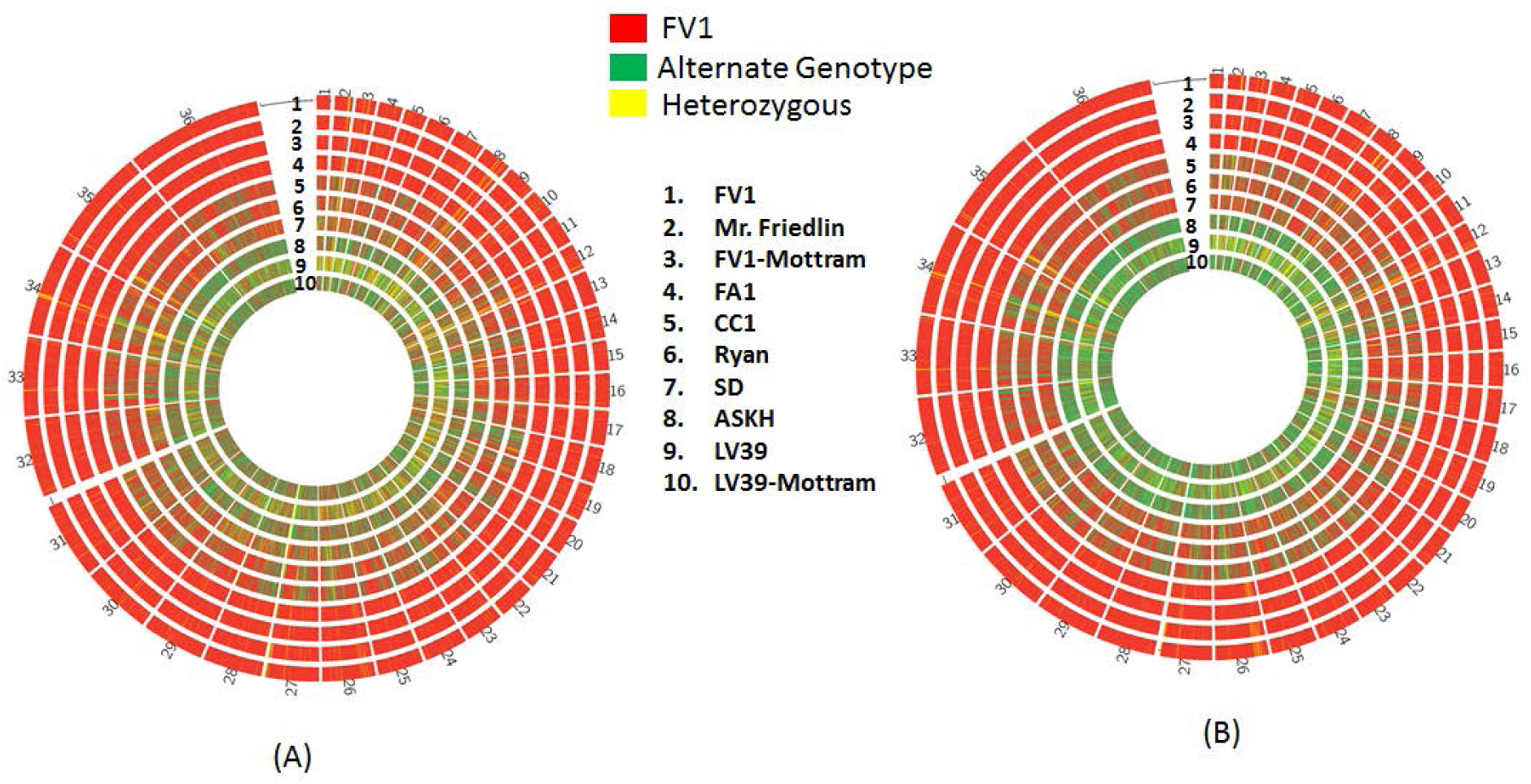
Allele compositions of *L. major* Strains. A) The *L. major* strains contain two major genotypes as indicated in red and green. The strains FV1 and FA1 are predominantly red with extremely low SNP counts and have identical haplotypes. The strains LV39 and ASKH on the other hand contain a SNP every ~500 bp, the majority of which are homozygous. The strains CC1, Ryan and SD are predominantly red genotype and have acquired mutations which are sparse but seem to occur in common blocks indicating a common origin of divergence from the red genotype. The SD strain is similar to CC1 and Ryan except that it has accumulated more mutations as evident from chromosomes 32 and 33. B) We eliminated private SNPs and redrew the allele compositions. The SD strain now looks similar to CC1 and Ryan suggesting their common origin and diversification that occurred due to independent mutations.

### The L. infantum Strains which were the products of extreme inbreeding show patterns similar to L. major

The patterns involving blocks of two different genotypes as seen in *L. major* are also observed in *L. tropica* groups 1 and 2. Similar patterns involving strains from Turkey were argued to be products of extreme inbreeding in *L. infantum* (Rogers et al. 2014b). Although not shown, the alternate genotypes in these *L. infantum* strains were assumed to be from *L. donovani*. We reanalyzed this data by including two *L. donovani* strains (Supplementary File4) to confirm if the blocks of alternate genotype were indeed from *L. donovani*. We used 157014 markers from the *L. donovani* strain *BPK035A1* that were homozygous and different from *L. infantum* strain *JPCM5*. The genotypes that emerged suggested that the strains from Turkey included blocks of two genotypes involving *L. infantum* and *L. donovani* (supplementary Figure 3). We performed simulations to verify if these patterns can be generated by extreme inbreeding by considering three different scenarios for mating and concluded that either successive matings of the outcrosses with one of the parental strains randomly or mating involving random selection of individuals from a population can generate these patterns (Supplementary File6)

**Supplementary Figure 3.**
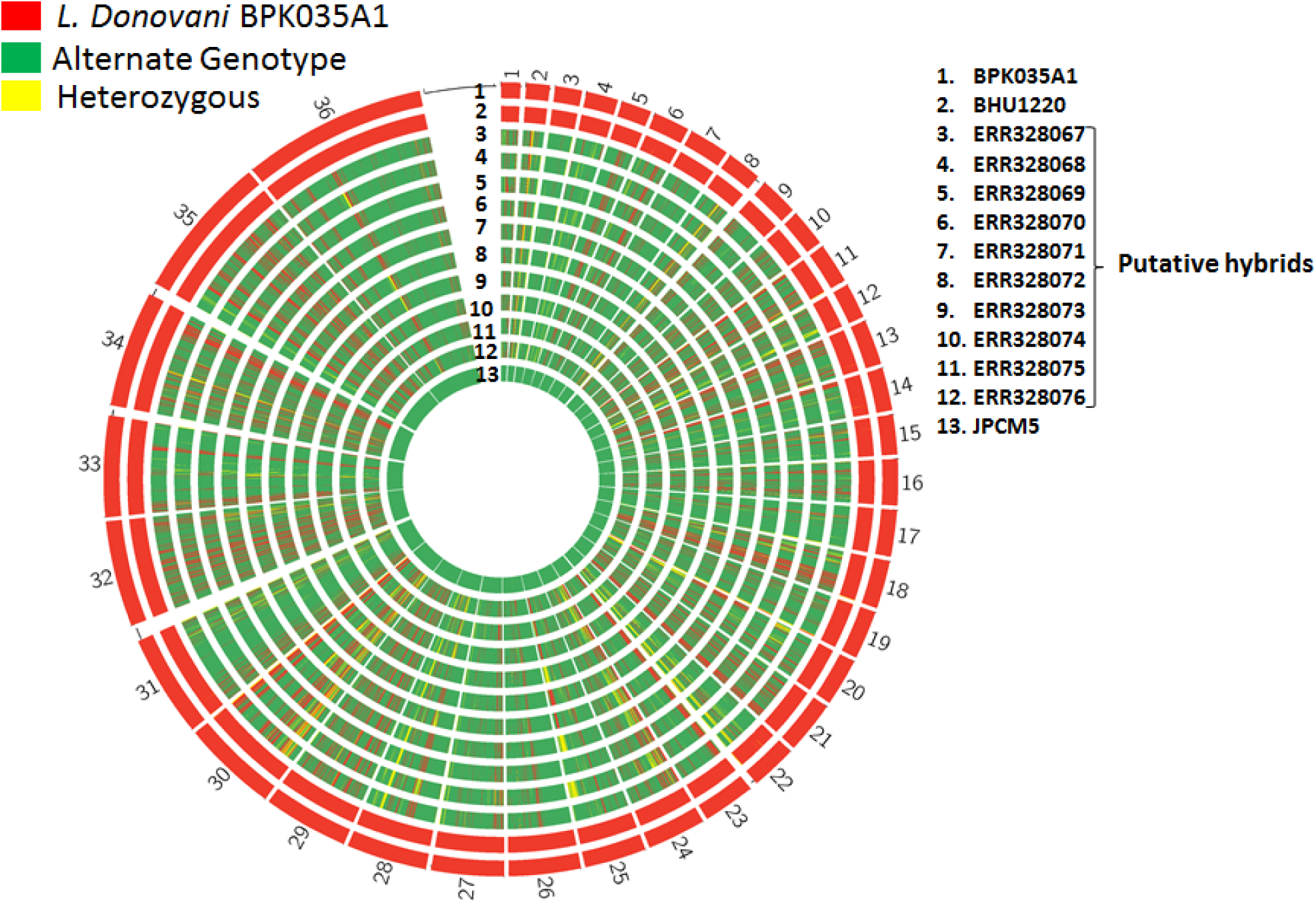
Reanalysis of *L. infantum* data previously argued to be hybrids. The SNPs where *L. donovani* strain *BPK035A1* (red) and *L. infantum* strain *JPCM5* (green) were homozygous and different were used for reanalysis. The patterns observed indicated that the putative hybrids belong to the *L. infantum* genotype but included blocks inherited from *L. donovani* suggesting their hybrid nature. The F1 hybrid generated between *L. infantum* and *L. donovani* perhaps underwent extreme inbreeding resulting in loss of heterozygosity.

## Discussion

Parasites of the genus *Leishmania* are responsible for a broad spectrum of human diseases, a diversity that is also seen in other aspects of their biology including their morphology, vector usage and the range of species the parasite is able to infect. This diversity is juxtaposed with the surprisingly high level of genetic conservation between strains and species. At the genetic level, this diversity could be explained by a small number of species-specific genes, epigenomic control or the presence of aneuploidy, however there have been challenges in studying the latter due to limitations with software tools often designed for diploid organisms. The aim of this study was to capture the genetic variation present in aneuploid genomes in the genus *Leishmania* and to understand the role that sex plays in driving diversity using a novel software program, POPSICLE, that we have adapted to analyze aneuploid organisms.

We began by employing traditional methods, including phylogenetic analysis and PCA, of publicly available WGS data for *Leishmania* and found the classifications to agree with subgenus, species and strain assignments. The two noted exceptions were later confirmed to be hybrids or coinfections (Supplementary Figure2). Aneuploidy, particularly mosaic aneuploidy is now considered to be an essential feature of *Leishmania* and it is hypothesized that *Leishmania* leverages genome plasticity as a means for survival and proliferation (Sterkers et al. 2012; Sterkers et al. 2014). We were therefore interested to know whether aneuploidy also correlated with the species assignments. While most chromosomes were diploid, chromosome 31 (chromosome 30 in *L. mexicana*) consistently contained more than 3 copies in all the species and strains that we studied. Although aneuploidy was routinely seen, it appeared to be random and did not correlate with the subgenus or species classifications. We next enumerated SNPs and zygosity profiles to assess variety in *Leishmania* (Figures 3&4). We were intrigued by the fact that most species were either predominantly homozygous or heterozygous and showed long runs of respective zygosities. We also observed that while some strains agreed between somies and allele frequency distributions, some did not (Figure 5). The strains that contained 90% or more homozygous SNPs or those that had low SNP counts had distributions that did not agree with the somies. This disagreement between the allele frequencies and somies could be due to mosaic aneuploidy and not due to allelic differences among the homologous chromosomes. The heterozygous lines on the other hand showed allele frequency distributions that matched the somies. They may also include heterozygous SNPs due to mosaic aneuploidy but in small numbers and were perhaps overshadowed by heterozygous SNPs that occur due to mating between two alternate genotypes. These clues raised the question of whether the strains with high heterozygous SNP counts were recent hybrids.

As traditional approaches such as phylogenetic analysis or dimensionality reduction are not capable of revealing shared ancestries, we constructed the population structure of *Leishmania* using our novel population genetics software, POPSICLE (Shaik et al. 2018), which we had upgraded for aneuploid genomes. The POPSICLE analysis showed that shared ancestries were present among closely related species such as *L. braziliensis* and *L. peruviana* (Figure 6). POPSICLE was particularly useful in helping decode why some strains labelled as one species showed similarities with other species. We performed additional analyses based on the clues provided by POPSICLE and found that MHOM ET1994, originally classified as *L. aethiopica*, was a hybrid between *L. aethiopia* and *L. tropica*. This confirms permissibility of more than one species in the sand fly vectors and the generation of inter-species hybrids in wild, reinforcing sex as one of the factors for speciation.

We applied POPSICLE specifically to *L. tropica* and found that the strains contained in our data set were categorized into four groups that correlated with their geography and were products of two distinct homozygous genotypes from Jordan (Group 1) and Palestine (Group 2). The strains from Afghanistan and India were mostly F1 hybrids, some of which contained a few homozygous segments that perhaps occurred due to LOH that happens during mating in *Leishmania* as has been observed in experimental hybrids (Inbar et al. 2013; Romano et al. 2014). In contrast, the heterozygous strains closer to Palestine and Jordan were a product of multiple cycles of mating. The predominant presence of F1 heterozygous strains in Afghanistan and India may be due to the heterozygous strains being sterile to each other. If they had mated, we would have expected to observe a greater variety in genotypes, as observed in strains assigned to Group 4 (Figure 8). The *DBKM* strain was an exception to this and was found to be a product of multiple cycles of mating, largely with strains from Group 1. We have eliminated the possibility of mating between heterozygous lines to generate the genotype of this stain due to the lack of genotypic variety expected from such a cross. This suggests that the strains present in Afghanistan and India are perhaps only capable of mating with homozygous strains as observed for DBKM, suggesting that it may have been a recent migrant to India. F1 hybrids that migrated to Afghanistan and India at an earlier time would have less chance to mate with the homozygous strains found in Palestine and Jordan due to geographic barriers. The heterozygous strains from Group 4, which are close to Palestine and Jordan, may therefore be a consequence of multiple rounds of sex with homozygous strains from Groups 1 and 2 but not with each other. Evidence of sex in this species suggests that sex is more common than previously argued and might be playing a key role in evolution of *Leishmania*.

The *L. tropica* strains from Groups 1 and 2, although were predominantly of one genotype, contained blocks of the alternate genotype (Figure 8). Genotypic profiles such as these were previously argued to be products of extreme inbreeding in *L. infantum* from Turkey even though the blocks of alternate genotype were speculated and not verified to be from a closely related species *<u>L. donovani</u>* (Rogers et al. 2014b). We re-analyzed this data by including a couple of *L. donovani* strains and concluded that the blocks of alternate genotype were indeed from *L. donovani*. We observed similar patterns in *L. major* which also contains two alternate genotypes and were largely of one of the two genotypes with blocks of SNPs from the alternate genotype suggesting they might also be products of extreme inbreeding. The lack of heterozygous strains such as those observed in *L. tropica* in these species may be an indication of the absence of recent hybridizations or due to sample size employed for studying these species. Simulated data that included two different genotypes and outcrosses generated using meiosis for several generations yielded intermittent blocks of two genotypes with little to no heterozygosity (Supplementary File 6). It is possible that *Leishmania* undergoes meiosis during mating, however the exact sexual strategy employed, and the frequency thereof, is unknown currently. Another possible explanation for the presence of alternate genotypes is lateral gene transfers, which were reported to be occurring in approximately 3% of genes (E.Vikeved et al. 2016). However, the proportion of genome with alternate genotype blocks observed in datasets we studied argues against this possibility. Complementary to the ancestries derived by POPSICLE, the dichotomy in zygosities may help to distinguish strains with recent hybridizations from those with multiple mating cycles.

Overall, the further development of POPSICLE to support the analysis of aneuploid genomes, such as those found in *Leishmania*, provides a unique tool for determining the shared ancestry of strains in this genus. POPSICLE helps to delineate the genetic relationships between strains both at the whole genome and individual chromosome level and is unique in its ability to distinguish hybrids from coinfections. When combined with the data provided by traditional population genetics analyses, POPSICLE provides new insights into the sexual strategies used by *Leishmania*. The proposed tool has broad applications to other genomes that exhibit mosaic aneuploidy such as Autism, Schizophrenia, Auto-immune diseases, Alzheimer’s, Cancers and *Fungi.*

## Methods

### POPSICLE employs a unique coding scheme to support aneuploid genomes

POPSICLE employs a unique coding scheme that leverages allele frequencies to determine ancestries. It is particularly useful for genomes with mosaic aneuploidy where heterogeneity in somies and haplotypes is seen (Figure 11A). These subgroups can independently acquire mutations which manifest as heterozygous alleles that do not match the average somy (Figure 11B), as they occur due to differences among populations and not due to allelic differences among the homologous chromosomes. To address this issue, POPSICLE accommodates allele frequencies by using a three-dimensional matrix of markers (M), samples (S) and nucleotides (4: corresponding to A, T, C and G). For example, 0:0.5:0.5:0 means that at a given marker, two nucleotides T and C are each present at 50% frequency. For most populations, there might only be two primary haplotypes but multiple copies of each of them. But, we have coded the data as MxSx4 to accommodate mutations independently acquired by haplotypes and to capture heterogeneity in the clonal population (Figure 11C). The strains are compared against the reference sequence and the markers in concordance are set to 0 and the rest are scored based on their deviation from the reference. POPSICLE groups the resulting numeric data for various “K” using K-means algorithm. The choice of optimum “K” is determined by employing the cluster evaluation metrics such as Dunn index (Dunn 1973; Davies and Bouldin 1979; Rousseeuw 1987). After fixing the value of “K”, POPSICLE runs the ancestry determination algorithm just once to facilitate better use of computational resources and therefore reducing the computational time.

**Figure 11.**
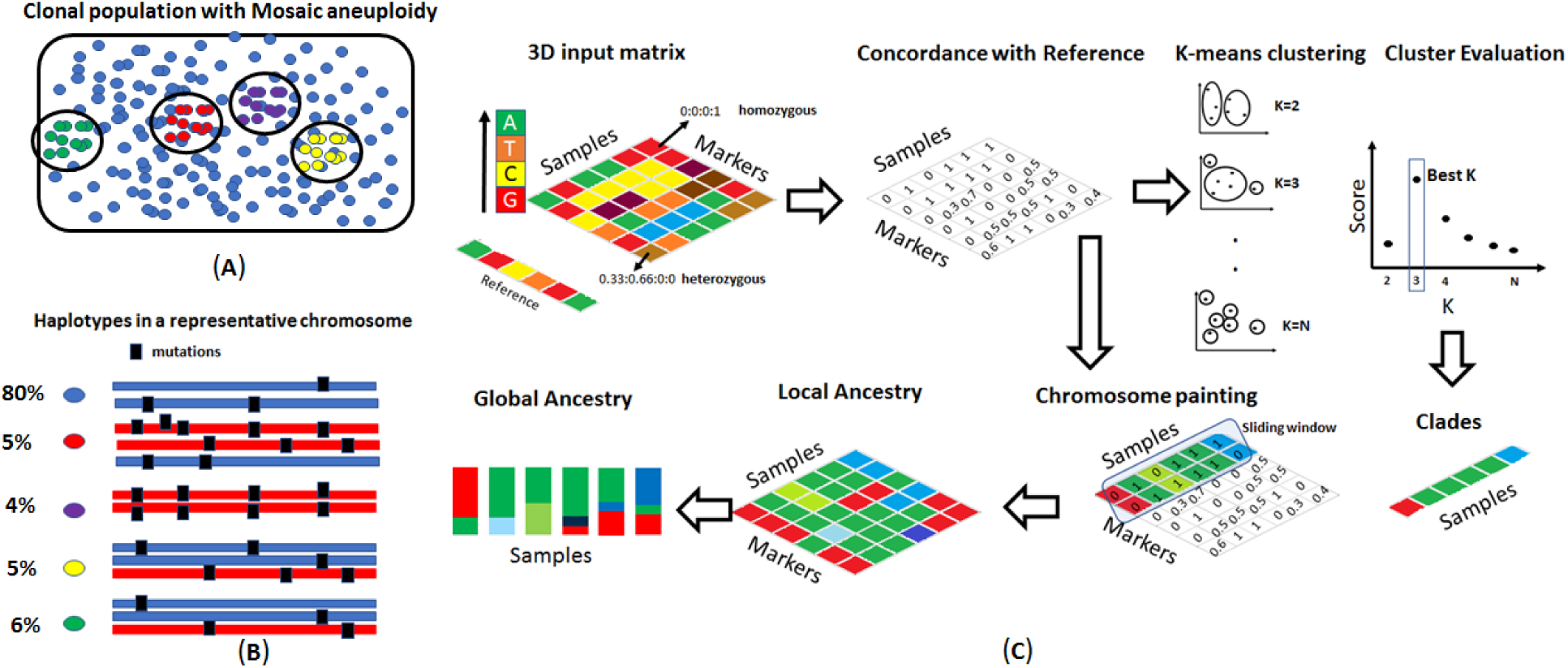
POPSICLE pipeline for population genetics in aneuploid genomes. A) In mosaic aneuploidy, certain subgroups of cells within a clonal population can independently develop aneuploidies and encompass unique haplotypes. B) Haplotypes in a representative chromosome show the aneuploidies and mutations acquired independently in each sub-population. Depending on the population size of these subgroups, aneuploidies and independent mutations acquired, the heterozygous loci may have allele frequencies that are different from overall ploidy of the organism. C) POPSICLE instead leverages allele frequencies to detect admixtures. POPSICLE compares these allele frequencies against a reference genome and finds genetic distances from the reference genome at each marker. The samples are subsequently clustered into groups (K) of 2, 3 to “N”, a user defined value. For various “K”, the clusters are evaluated for intra-cluster compactness and inter-cluster separation using metrics such as Dunn-Index and Davies-Bouldin index and “K” value that offers the best value are selected automatically. The markers within a sliding window across the genome are independently clustered and compared against the original clade assignments. The strains are assigned locally to the existing clades or new clades depending on the similarities of the strains with the original clades. This reveals regions with shared ancestries, new ancestries not included in the cohort and the differences across the genome. These genome-wide ancestries are summarized to generate global ancestries.

**Supplementary Figure 4.**
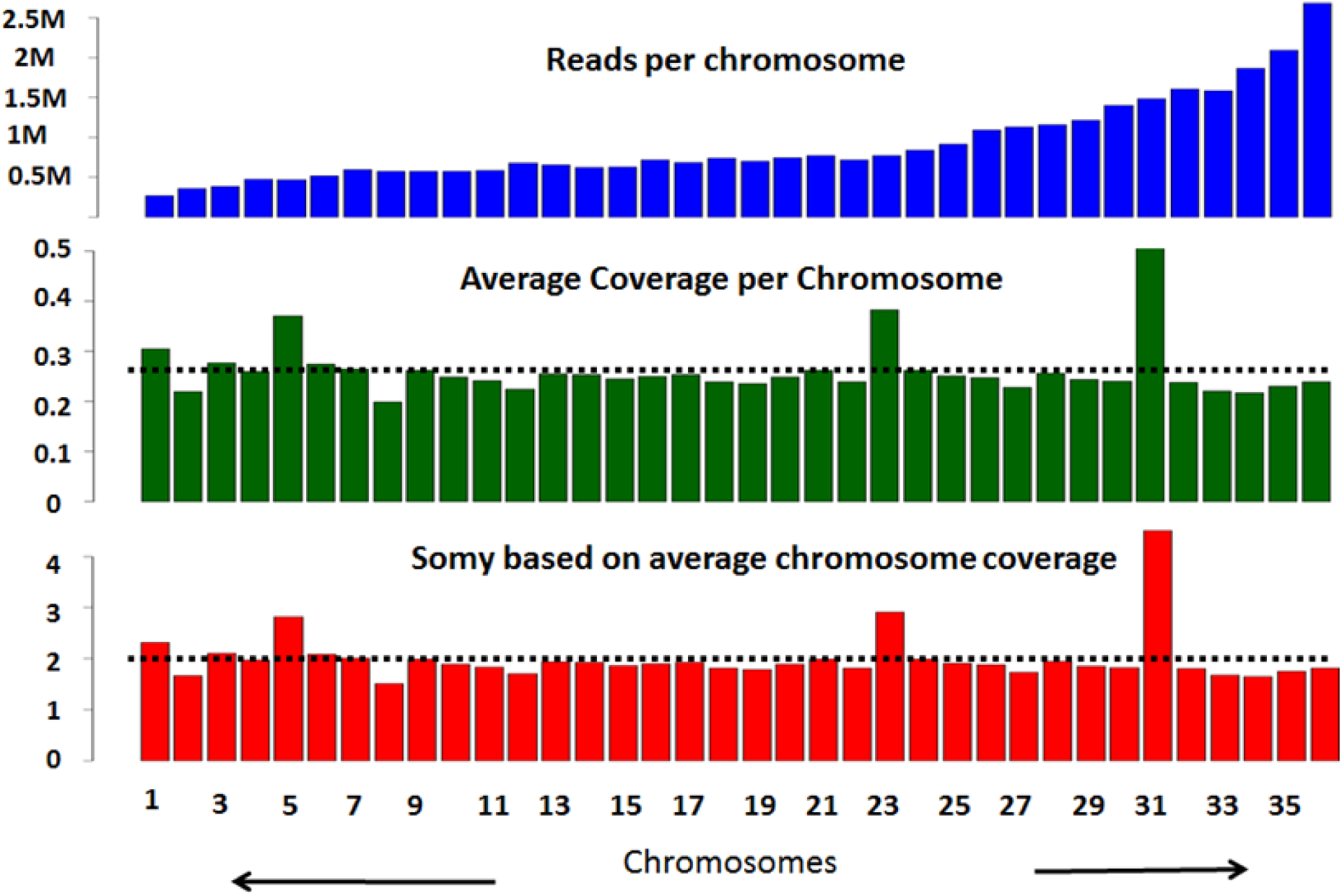
The reads aligned per chromosome depend on the somy of the chromosome and length of the chromosome. The reads per chromosome are normalized using chromosome lengths, normalized to average coverage of the genome and are subsequently scaled to the ploidy of the organism to obtain somies

### POPSICLE determines global and local ancestries using chromosome-painting like approach

POPSICLE divides the genome into non-overlapping sliding windows of user defined size (e.g. 1kb) and assigns the local profiles to the existing clades or the new clades as needed. This improvement is significant because unlike the existing algorithms, POPSICLE does not force the association of haploblocks to predefined number of clusters. The clusters are collapsed into one if the strains globally associated to multiple clades are similar locally or new clades are created to accommodate new populations (Figure 11E). These new clades are represented using colors other than the colors associated with the original clades in POPSICLE. The local ancestry determination is repeated by sliding the window across the genome (Figure 11F). The local ancestry patterns reveal genomic segments that are shared, unique segments that are acquired from new populations and the overall genomic divergence. The proportion of different populations present in each strain can be summarized to reveal global ancestries.

### Other Bioinformatic methods

The short reads from the Illumina sequencer (100bp) were aligned to the genome of interest using BWA using default parameters (Li et al. 2009). Reads aligned per chromosome are a consequence of somy of chromosome and the chromosome length. We nullified the effect of chromosome length by diving the read counts per chromosome by chromosome lengths. Since most chromosomes are expected to be disomic, we subsequently normalized these values using average coverage across the genome. The normalized values were scaled to the ploidy of the organism to reveal somies (Supplementary Figure 4). Single nucleotide polymorphisms were obtained using samtools, mpileup function and using the –ploidy-file feature, which calls SNPs by taking chromosomal somies into account (Li et al. 2009).

SNP densities across the genome were obtained in sliding non-overlapping windows of 5kb and plotted as line plots in circos V. 0.69 (Krzywinski et al. 2009). The proportions of heterozygous and homozygous SNPs were also counted in sliding blocks of 5kb and plotted as histogram plots in circos where colors for the histogram bars were assigned based on zygosity of the block. If the block was 80% or more homozygous, a blue color was assigned, if the block was 80% or more heterozygous, a red color was assigned, and yellow color was assigned otherwise.

Allele frequency plots were generated by employing the heterozygous SNPs. Read alignments to loci containing heterozygous SNPs will have some reads representing the reference allele and some others representing the alternate allele. The proportion of reads representing each allele type were obtained for all heterozygous SNPs. A histogram of these allele frequencies was depicted to yield allele frequency distributions. For disomic chromosomes, all heterozygous alleles will have 50% of reads representing the reference allele and the other 50% representing the alternate allele. Although minor variations are expected due to bioinformatics error, a peak is expected at 0.5. Similarly, depending on the somy of the chromosome, peaks are observed at different allele frequencies. When the heterozygosity is mainly a consequence of aneuploidy and not due to differences among haplotypes, the correlations between somies and allele frequency distributions are not seen.

Synteny map was generated by creating a nucleotide blast database of LmjF33 genome and performing a nucleotide blast of *L. braziliensis M2904* V.33 genome against that database. All blast hits with identity of 70% or more, e-value less than 0.001 and minimum segment size of 500bp were drawn using ribbon links in circos.

## Acknowledgements

The author would like to thank Andrea Paun and Asis Khan for critical reading and editing of the manuscript. This study used the Office of Cyber Infrastructure and Computational Biology (OCICB) High Performance Computing (HPC) cluster at the National Institute of Allergy and Infectious Diseases (NIAID). The authors are supported by the Intramural Research Program of the NIAID at the National Institutes of Health, Bethesda, MD.

